# Benefits and costs of a hypercapsule and the mechanism of its loss in a clinical isolate of *Acinetobacter baumannii*

**DOI:** 10.1101/2025.08.04.668531

**Authors:** Chaogetu Saren, Ken-Ichi Oinuma, Taishi Tsubouchi, Arata Sakiyama, Masato Suzuki, Mamiko Niki, Yukihiro Kaneko

**Author notes:** Address correspondence to Ken-Ichi Oinuma.

## Abstract

*Acinetobacter baumannii* is an opportunistic pathogen in which capsule production is closely linked to immune evasion and environmental persistence. Recent studies have described two seemingly contradictory phenomena—increasing prevalence of capsule-overproducing clinical isolates and frequent isolation of capsule-deficient variants. The biological significance of these phenomena remains unclear. In this study, we analyzed a clinical isolate, OCU_Ac16b, which spontaneously gives rise to two phenotypically distinct variants: the L type forming large colonies with an extremely thick capsule, and the S type forming small colonies with a substantially reduced or absent capsule. When L-type cells were cultured in test tubes under low-shaking conditions, S-type variants reproducibly emerged, constituting 45–80% of the population within 24 hours. Whole-genome sequencing revealed that this conversion is driven by distinct mutations in the capsular polysaccharide synthesis cluster, including insertion sequence (IS) insertions and a single-nucleotide deletion. PCR analysis confirmed that IS insertions in *wzy* (polysaccharide polymerase) occur within individual L-type colonies prior to liquid culture, providing a likely explanation for the rapid and reproducible emergence of S-type variants. Phenotypic characterization demonstrated a biological trade-off, with L-type cells exhibiting enhanced resistance to serum killing, desiccation, and certain β-lactam antibiotics, whereas S-type cells showed superior surface attachment, increased biofilm formation, and a growth advantage under oxygen-limited conditions. Our findings uncover a highly reproducible, mutation-driven capsule switching mechanism that enables rapid phenotypic adaptation to changing environments. This phenotypic heterogeneity has significant implications for pathogenesis, persistence, diagnostic evaluation, and clinical management.

**Importance:** *Acinetobacter baumannii* is a clinically important opportunistic pathogen that exhibits striking phenotypic diversity. In particular, some clinical isolates produce unusually thick capsules, which are thought to contribute to immune evasion and persistence, while others lack capsule altogether. However, the biological significance of these contrasting phenotypes has remained unclear. In this study, we analyzed a clinical isolate that spontaneously gives rise to capsule-deficient variants from a hypercapsulated form. We found that the conversion is driven by spontaneous mutations in capsule biosynthesis genes that arise naturally within colonies, while the expansion of capsule-deficient cells is promoted under oxygen-limited conditions. The two variants differed in serum resistance, desiccation tolerance, growth characteristics, and antibiotic responses, revealing a trade-off between protective barriers and environmental adaptability. These findings provide new insights into how *A. baumannii* balances survival strategies through genetic and phenotypic heterogeneity, with important implications for diagnosis, treatment, and bacterial persistence in clinical settings.

## Introduction

*Acinetobacter baumannii* is an opportunistic Gram-negative pathogen that has emerged as a major cause of hospital-acquired infections worldwide. *A. baumannii* infections commonly occur in immunocompromised patients and are associated with ventilator-associated pneumonia, bloodstream infections, wound infections, and urinary tract infections (1–3). A defining characteristic of *A. baumannii* is its ability to rapidly acquire antimicrobial resistance through a wide array of mechanisms, including horizontal gene transfer via plasmids and natural transformation (1, 2, 4, 5). The genome of *A. baumannii* harbors numerous insertion sequences (ISs), which are minimal transposable elements encoding transposase. ISs often enhance the expression of chromosomal resistance genes, such as *bla*_OXA-51_, by inserting strong promoter sequences (6, 7). In addition, ISs can disrupt various genes critical for antimicrobial susceptibility, such as those encoding outer membrane porins (7, 8).

Beyond resistance, *A. baumannii* exhibits remarkable adaptability to diverse environments, in part facilitated by biofilm formation and capsule production (9). Capsules have been shown to contribute to immune evasion, disinfectant resistance, and desiccation tolerance, making them a significant determinant of bacterial pathogenicity (10). A recent study reported that an increasing number of modern clinical isolates of *A. baumannii* display thicker capsules and higher levels of capsulation compared to historically established strains (11). Other studies described the rising prevalence of mucoid *A. baumannii* isolates, which exhibit thicker capsules and are frequently associated with long-term infections or heightened virulence (12, 13). However, the underlying mechanisms and biological consequences of this phenotypic shift remain largely unexplored.

We recently reported on the clinical isolation and genomic characterization of *A. baumannii* OCU_Ac16a, harboring *bla*_NDM-1_, *bla*_TMB-1_, and *bla*_OXA-58_, and its clone, OCU_Ac16b, which lacks *bla*_NDM-1_ but retains *bla*_TMB-1_ and *bla*_OXA-58_ (14). While our initial focus was on their antimicrobial resistance, subsequent observations made during the course of this study revealed that OCU_Ac16b exhibits notable variations in colony size and capsular morphology. More specifically, a subpopulation of OCU_Ac16b exhibited a remarkably thick capsule (hypercapsule), forming relatively large colonies on agar plates (designated the L type), while other variants with either no capsule or a much thinner one formed smaller colonies (designated the S type). Moreover, we found that S-type variants emerged within the L-type population during liquid culture, replacing 40–80% of the population within 24 hours. This phenomenon was consistently observed in repeated independent experiments, with near-complete reproducibility.

In this study, we aimed to elucidate the clinical relevance and mechanistic basis of this L-to-S phenotypic conversion in OCU_Ac16b. Through genome sequencing of representative L- and S-type variants, we found that the conversion is driven by distinct mutations, such as multiple IS insertions and at least one single-nucleotide deletion, within the capsular polysaccharide synthesis (*cps*) cluster. PCR screening of individual L-type colonies revealed that some of those mutations, specifically insertions of an IS into a polysaccharide polymerase gene *wzy*, were already present within many colonies prior to liquid culture, although likely limited to a small subpopulation of cells. These findings highlight a previously unrecognized phenotypic switch in *A. baumannii*, driven by frequent mutations arising at multiple sites in the *cps* cluster that lead to capsule loss. Our findings also shed light on the functional trade-offs associated with hypercapsule production: while L-type cells exhibit enhanced resistance to serum killing, desiccation, and certain β-lactam antibiotics, S-type variants demonstrate superior surface attachment, biofilm formation capacity, and a fitness advantage under oxygen-limited conditions. By demonstrating that these genetic changes confer distinct advantages and disadvantages under different environmental conditions, this study illustrates the dual nature of hypercapsule formation as both a protective and burdensome trait. Our study underscores the importance of considering dynamic and heterogeneous bacterial subpopulations when evaluating virulence, resistance, and treatment outcomes.

## Results

### Discovery and characterization of large colony and small colony variants of OCU_Ac16b

To investigate the phenotypic heterogeneity of OCU_Ac16b, we selected representative L-type and S-type variants (designated L1 and S1, respectively) and compared their colony morphology and capsule production. On cation-adjusted Mueller Hinton (CAMH) agar plates, L-type colonies appeared approximately 1.5 times larger and more opaque than S-type colonies (Fig. 1A). Light microscopy with India ink-negative staining revealed a prominent hypercapsule surrounding L-type cells, whereas no visible capsule was observed in S-type cells (Fig. 1B).

**Fig. 1.**
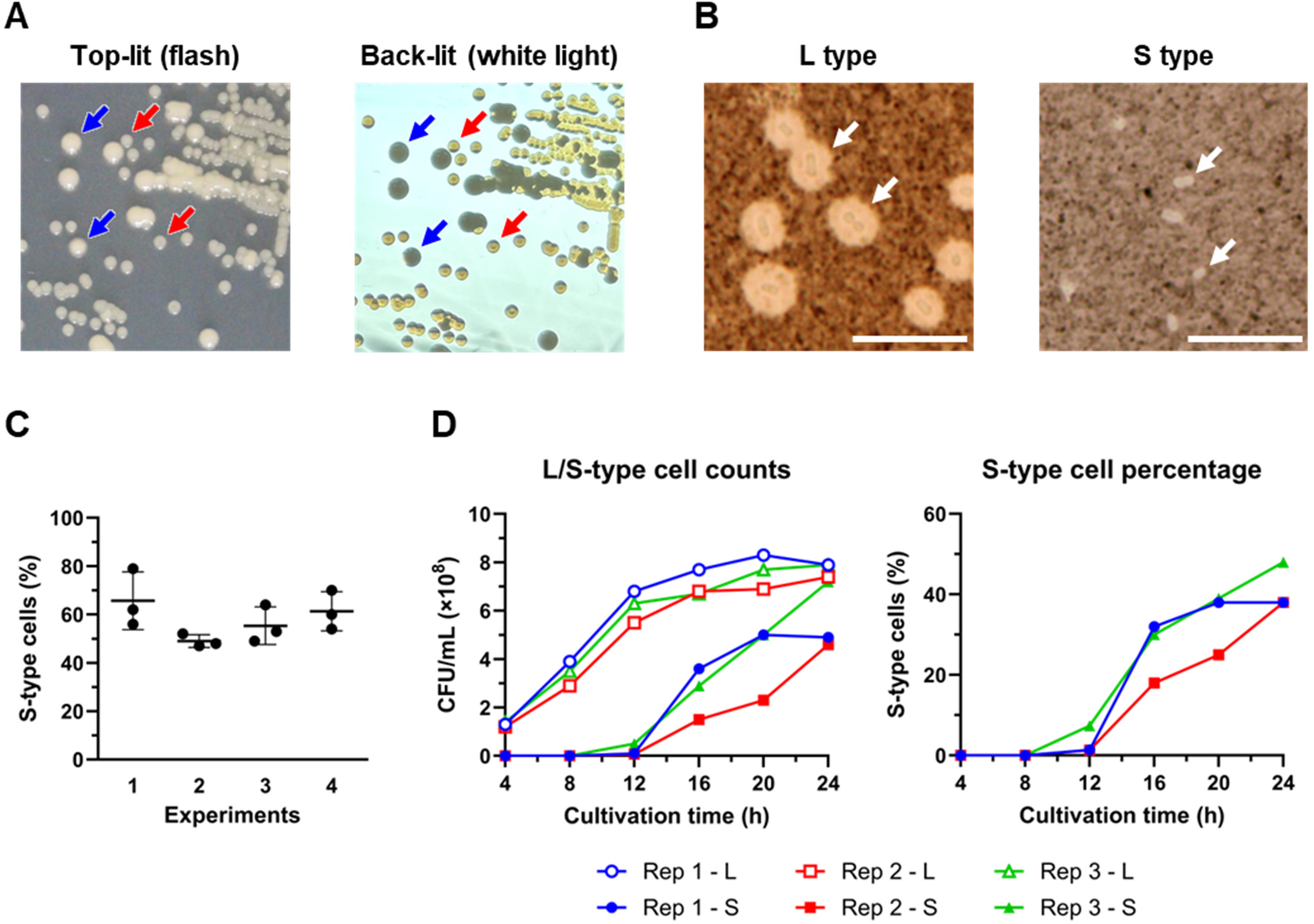
Discovery and characterization of large-colony (L-type) and small-colony (S-type) variants of *Acinetobacter baumannii* OCU_Ac16b. (A) OCU_Ac16b forming both L-type and S-type colonies was grown on cation-adjusted Mueller Hinton (CAMH) agar plates overnight at 37°C. The left panel shows colonies photographed from above with flash illumination. The right panel shows colonies photographed with back illumination; the plate was held approximately 50 cm in front of a white light source. Representative L-type (blue arrows) and S-type (red arrows) colonies are indicated. (B) Negative staining with India ink of bacterial cells from L-type and S-type colonies. Cells were observed under a light microscope. L-type cells exhibited a prominent halo, indicating the presence of an extremely thick capsule (hypercapsule), whereas S-type cells showed no visible capsule. Arrows indicate representative cells. Scale bar, 10 μm. (C) Conversion of L-type to S-type variants during liquid culture. Single colonies of the L-type strain (L1) were inoculated into 3 mL of CAMH broth in 14-mL test tubes and incubated overnight at 37°C with shaking at 120 rpm. Cultures were then plated onto CAMH agar plates to evaluate the colony morphologies. Four independent experiments were performed, each using three separate colonies. (D) Time-course of S-type cell emergence from L-type cultures. Three independent L1 cultures (Rep 1 to 3) were grown in CAMH broth (3 mL, 37°C, shaking at 120 rpm), and samples were taken at indicated time points to determine CFUs of L-type and S-type cells. The left graph shows the CFUs for each type, and the right graph indicates the percentage of S-type cells in the population.

During liquid culture, S-type colonies reproducibly emerged from L-type populations. When single L-type colonies were inoculated into 3 mL of CAMH broth in 14-mL tubes and incubated at 37°C with shaking at 120–130 rpm for 24 hours, S-type cells comprised up to 50% of the population. This conversion was unidirectional, as S-type cultures did not yield L-type variants. The phenomenon was unlikely to be due to contamination, as S-type colonies were absent when L1 stocks were directly plated or when L-type colonies were transferred to agar without liquid culture. To assess reproducibility, we conducted four independent L-type culture experiments, each with triplicate cultures under standardized conditions (CAMH broth, 37°C, 120 rpm). S-type cells consistently emerged, comprising 47–79% of the population after 24 hours (Fig. 1C). Time-course analysis of three independent L1 cultures showed that S-type cells were not detected at 8 hours but became detectable in all cultures by 12 hours, reaching 38–48% by 24 hours (Fig. 1D). These results demonstrate reproducible L-to-S phenotypic conversion during liquid culture.

### Identification of genetic determinants of L-to-S phenotypic conversion

Genome comparison between L1 and S1 strains revealed an 875-bp chromosomal inversion at the *brnT*/*brnA* locus (chr: 3,645,941–3,646,815), which is presumed unrelated to capsule switching. More importantly, S1 harbored a 1,040-bp insertion sequence (IS*Aha2*) within the *wzy* gene encoding polysaccharide polymerase in the *cps* cluster (Fig. 2A, B). The insertion at positions 613–622 of *wzy* created a 9-bp direct repeat (TATTGGCGT) flanking the IS element, with the transposase oriented opposite to *wzy*. This *wzy* disruption likely accounts for the capsule loss in S1.

**Fig. 2.**
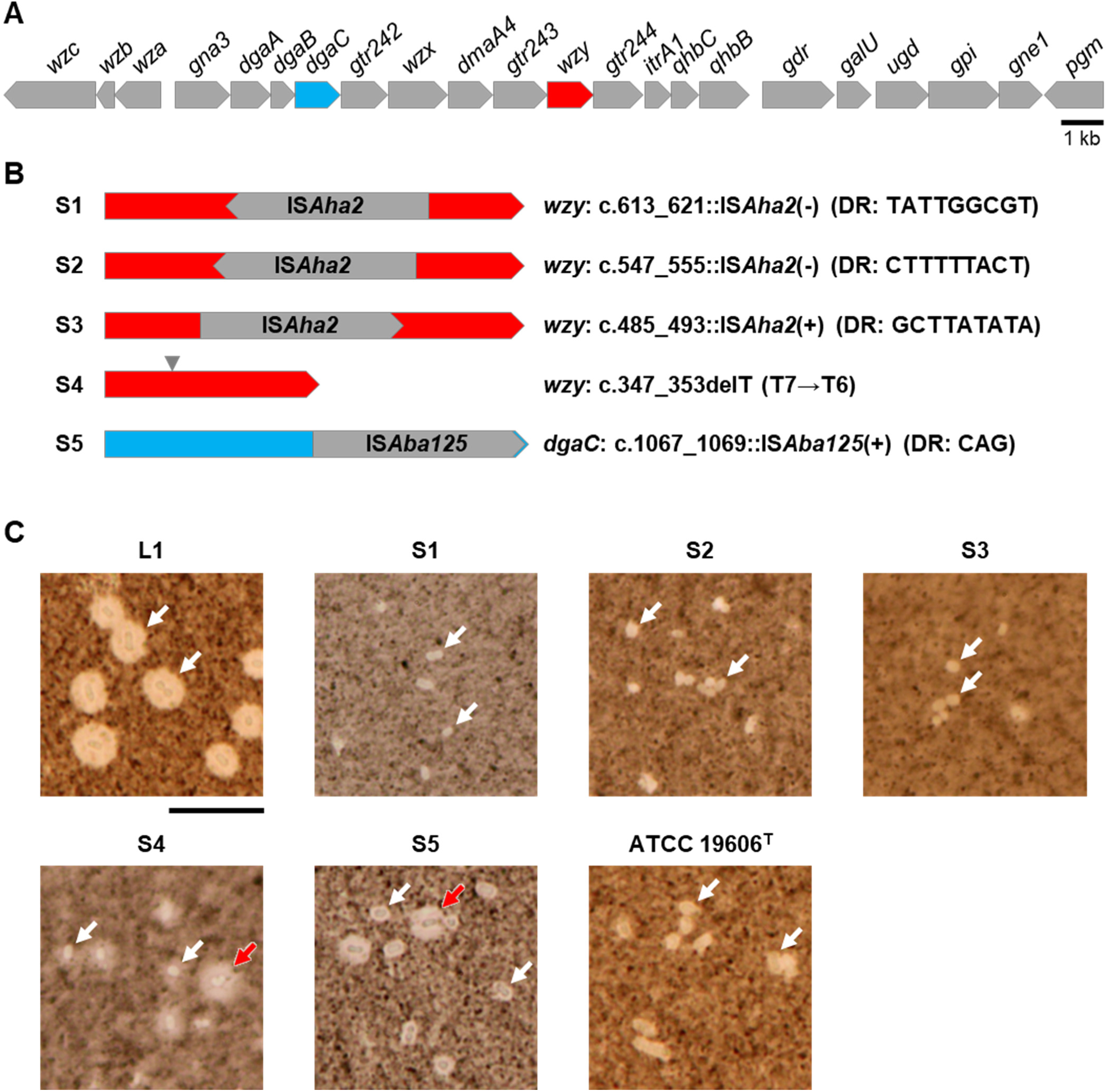
Identification of mutations responsible for the phenotypic conversion from L-type to S-type in *Acinetobacter baumannii* OCU_Ac16b. (A) Schematic representation of the entire capsule polysaccharide synthesis gene cluster in the L-type strain. *wzy* and *dgaC* are highlighted in red and light blue, respectively. (B) Schematic representation of genetic alterations identified in S-type variants S1–S5. S1–S4 harbor mutations in *wzy*, while S5 contains an insertion in *dgaC*. “c.” indicates the position within the coding sequence of the gene. (+) and (-) indicate that an insertion sequence is oriented in the same or opposite direction, respectively, relative to the target gene. DR, direct repeat sequence observed at both ends of the insertion elements. (C) Capsule phenotypes of L1 and five S-type variants (S1–S5), as well as the reference strain ATCC 19606^T^, visualized by negative staining with India ink and light microscopy. Images of L1 and S1 are reused from Fig. 1B. S4 and S5 exhibited a mixture of cells with varying capsule thickness, including both thinly and thickly encapsulated cells. Representative thinly encapsulated cells are indicated with white arrows, and thickly encapsulated cells with red arrows. Scale bar, 10 μm.

To assess the reproducibility of this mutation event, we isolated 16 S-type colonies newly emerged from an L-type culture and screened them by PCR for IS insertions in *wzy*, using primer pairs listed in Table 1. Four of the 16 strains showed PCR product sizes indicative of IS insertion. Subsequent sequencing of the putative IS-inserted PCR products revealed two additional IS*Aha2* insertions at distinct sites within *wzy*, found in strains S2 and S3 (Fig. 2B). We selected four representative strains for further analysis: S2 and S3, which carried IS*Aha2* insertions in *wzy*, and S4 and S5, which lacked such insertions. Microscopy confirmed that S2 and S3, like S1, lacked visible capsule structures, whereas S4 and S5 exhibited heterogeneous capsule formation, with both thickly and thinly encapsulated cells observed (Fig. 2C). Notably, the capsule in S5 appeared as a well-defined, sharply demarcated halo surrounding the cells, whereas that in S4 was often more diffuse and irregular, lacking a uniform layer around the cell surface. Genome sequencing revealed no additional mutations in S2 and S3 beyond the IS*Aha2* insertions in *wzy*. S4 had a single thymine base deletion in *wzy*, and S5 harbored an IS*Aba125* insertion in *dgaC*, a putative aminotransferase gene located within the *cps* cluster (Fig. 2B). These findings suggest that spontaneous mutations in key *cps* genes are the direct cause of L- to-S phenotypic conversion in most cases, if not all.

**Table 1.**
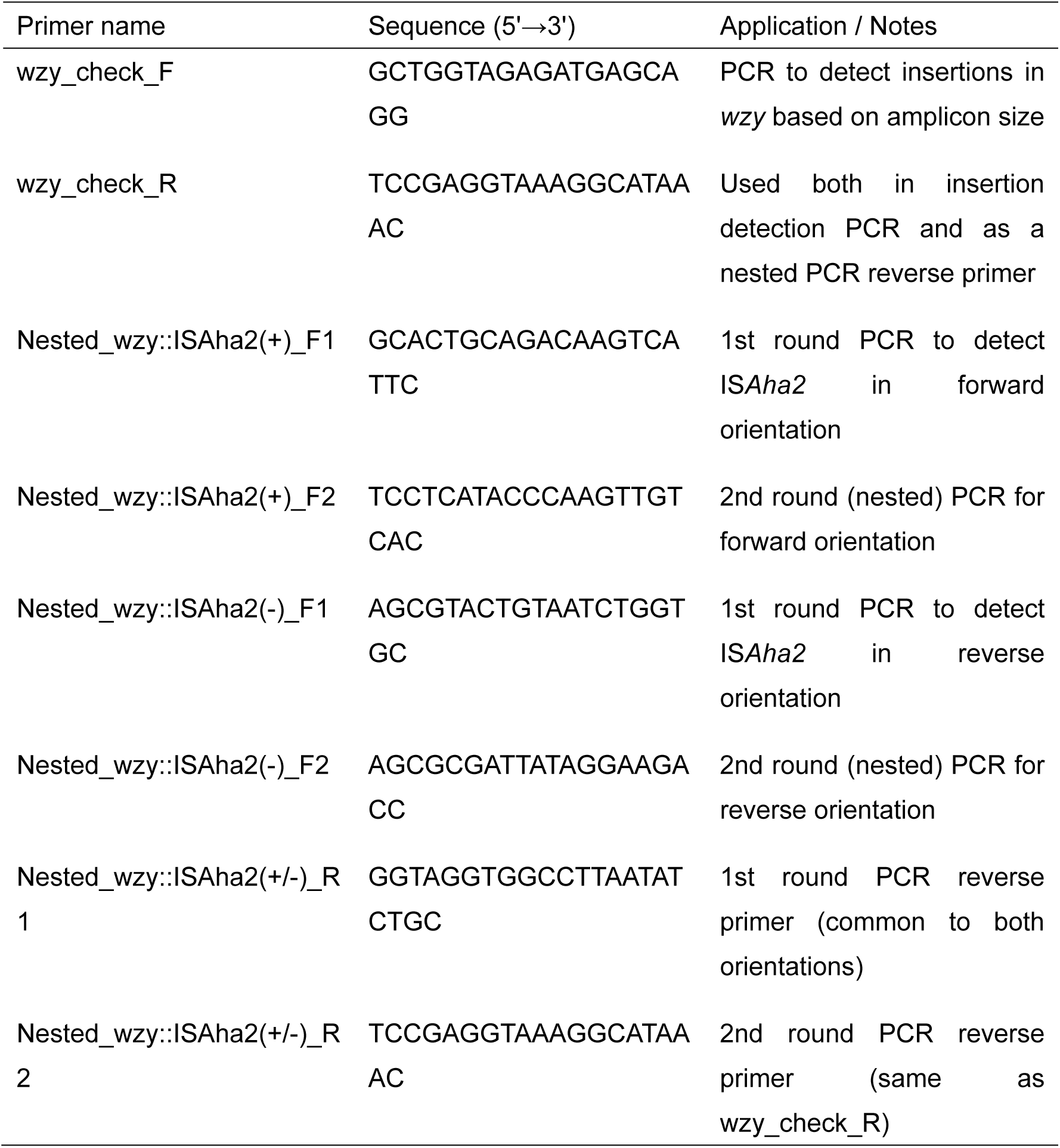
Primers used in this study.

### The clinical relevance of the L-to-S phenotypic conversion

To assess the clinical relevance of capsule switching, we compared the antimicrobial susceptibility, serum resistance, desiccation tolerance, and biofilm formation of L-type and S-type variants. Antimicrobial susceptibility testing revealed that L1 cells showed higher resistance to certain β-lactam antibiotics, with MICs twice as high for imipenem and more than four times higher for cefepime compared to S1 cells (Table 2). This suggests that the hypercapsule provides protective effects against specific β-lactams.

**Table 2.** Minimum inhibitory concentrations (MICs) of 12 antibiotics against *A. baumannii* strains.

| Strain | AMP | PIP | CAZ | FEP | IPM | MEM | KAN | AMK | LVX | MIN | TGC | CST |
| --- | --- | --- | --- | --- | --- | --- | --- | --- | --- | --- | --- | --- |
| ATCC 19606<sup>T</sup> | >256 | 16 | 4 | 8 | 0.25 | 0.25 | 8 | 16 | 0.25 | 0.064 | 1 | 0.094 |
| L1 | >256 | >256 | >256 | >256 | 16 | >32 | 4 | 4 | 0.5 | 0.064 | 0.25 | 0.125 |
| S1 | >256 | >256 | >256 | 64 | 8 | >32 | 4 | 4 | 0.5 | 0.047 | 0.25 | 0.094 |

MIC values are given in µg/mL. AMP, ampicillin; PIP, piperacillin; CAZ, ceftazidime; FEP, cefepime; IPM, imipenem; MEM, meropenem; KAN, kanamycin; AMK, amikacin; LVX, levofloxacin; MIN, minocycline; TGC, tigecycline; CST, colistin. All MICs were determined using Etest (bioMérieux).

To evaluate the impact of L-to-S phenotypic conversion on serum resistance, we tested the survival of L1, S1–S5, and ATCC 19606^T^ strains in pooled human serum at final concentrations of 2.5% and 10% (Fig. 3A). L1 exhibited high resistance, while S1–S5 showed substantially reduced survival, indicating that capsule loss or reduction generally compromises serum resistance. We note, however, that serum resistance varied among the S-type strains, with S5 showing relatively higher survival. This may suggest that capsule quality, in addition to quantity, influences serum resistance. Moreover, ATCC 19606^T^ also exhibited high serum resistance despite being capsule-deficient or possessing only a very thin capsule, implying the involvement of capsule-independent mechanisms, such as recruitment of complement regulatory factors by outer membrane proteins (15). These findings indicate that L-to-S conversion generally reduces serum resistance, although alternative mechanisms may provide protection in certain strains.

**Fig. 3.**
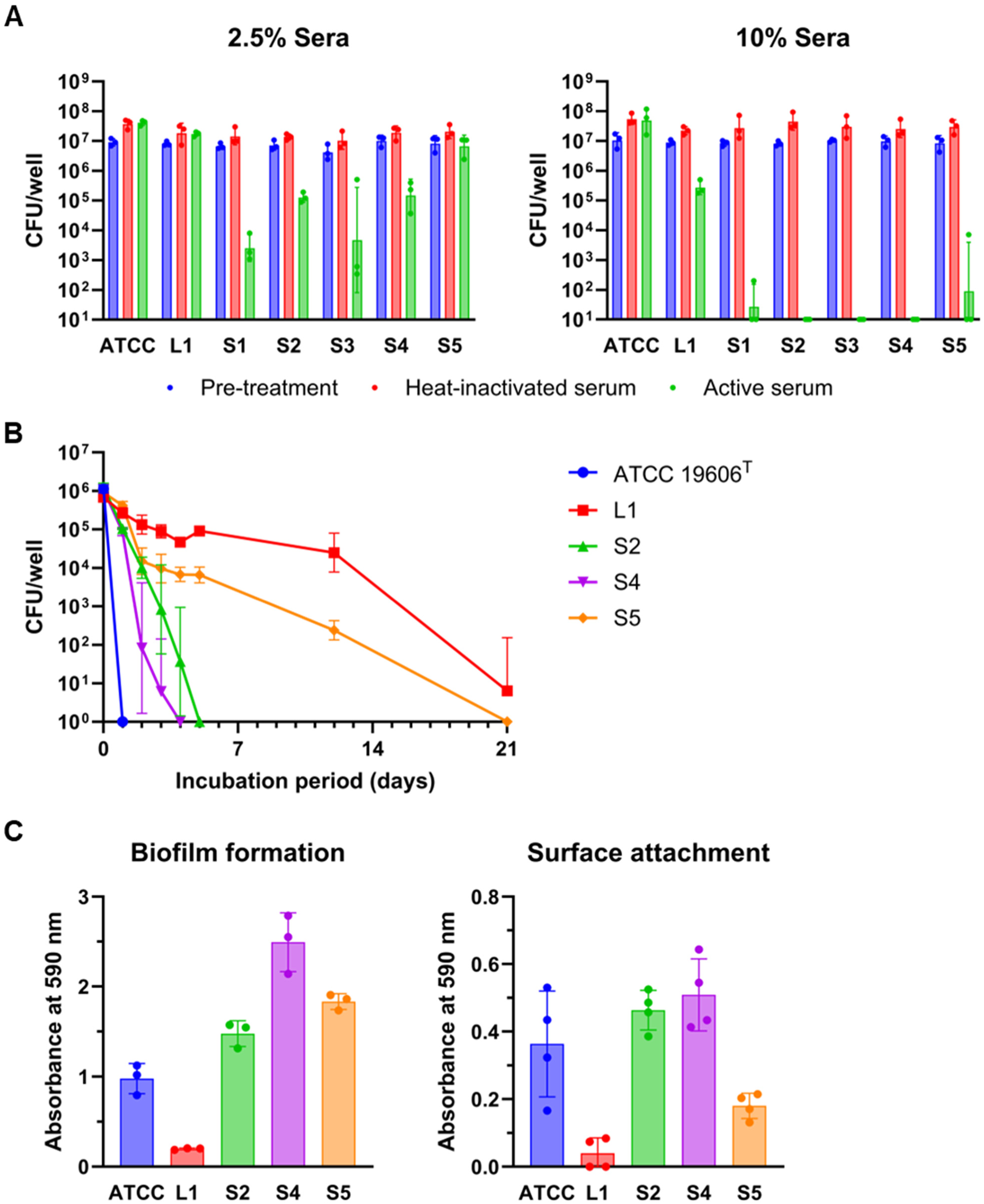
Phenotypic characterization of L-type and S-type variants reveals functional trade-offs associated with capsule production. (A) Serum resistance assay. Strains ATCC 19606^T^ (ATCC), L1, and S1–S5 were tested for survival in pooled human serum at final concentrations of 2.5% (left panel) and 10% (right panel). Cell suspensions were incubated in serum at 37°C with shaking at 100 rpm for 2 hours. Blue bars represent CFU prior to serum exposure and incubation, red bars represent CFU after incubation in heat-inactivated serum, and green bars represent CFU after incubation in active serum. Data represent geometric means ± geometric SD from three biological replicates (independent cultures initiated from single colonies), with individual data points shown. (B) Desiccation tolerance assay. Strains ATCC 19606^T^, L1, S2, S4, and S5 were tested for survival under desiccating conditions at 36°C and 13.1% (SD 2.8) relative humidity. Cell suspensions were spotted in 96-well plates and incubated under desiccating conditions. At designated time points (1, 2, 3, 4, 5, 12, or 21 days), cells from separate wells were rehydrated and plated to determine viable CFU. Data represent geometric means ± geometric SD from three independently prepared cultures, each derived from a different single colony. Values below the detection limit (1 CFU/mL) were plotted as 1 for visualization on a log scale. Lower error bars are not shown for L1 (day 21) and S4 (day 3) because they fall below the lower limit of the log-scaled Y axis (1 CFU/mL). (C) Biofilm formation and surface attachment assays. Strains ATCC 19606^T^, L1, S2, S4, and S5 were evaluated. Left panel: Biofilm formation was assessed after 48 hours of static cultivation in Brain Heart Infusion broth at 37°C in 24-well polystyrene plates. Right panel: Surface attachment was evaluated after 2 hours of cultivation in cation-adjusted Mueller Hinton broth at 37°C with shaking at 100 rpm. Both assays used crystal violet staining, and absorbance was measured at 590 nm. Data represent means ± SD from three or four biological replicates, with individual data points shown.

We next evaluated the desiccation tolerance of L1 and selected S-type variants (S2, S4, and S5), using ATCC 19606^T^ as a reference (Fig. 3B). L1 cells exhibited the highest tolerance, with viable cells detectable even after prolonged desiccation. In contrast, S-type variants showed markedly reduced survival, and ATCC 19606^T^ was the most sensitive. S5, which retained a capsule of heterogeneous thickness, showed intermediate tolerance. Interestingly, S4, despite also possessing a heterogeneous capsule, showed desiccation sensitivity comparable to or even greater than that of the non-encapsulated strain S2. These findings underscore the importance of the capsule in protecting against desiccation and suggest that not only its presence or thickness but also its structural integrity may influence the degree of protection.

We evaluated biofilm formation in L1 and three selected S-type variants (S2, S4, and S5), with ATCC 19606^T^ included for comparison. After 48 hours of static cultivation in Brain Heart Infusion (BHI) broth, all S-type strains produced more biofilm than L1, with levels ranging from 7.5- to 12.7-fold that of L1 (Fig. 3C, left). To investigate the underlying mechanism, we assessed surface attachment and found that S2, S4, S5, and ATCC 19606^T^ exhibited noticeably greater adherence than L1 (Fig. 3C, right), suggesting that enhanced attachment contributes to increased biofilm production. These findings underscore that L-to-S conversion may promote environmental persistence and influence infection outcomes by enhancing biofilm formation and surface colonization.

### S type has a fitness advantage under oxygen-limited conditions

Among the tested media, Tryptic Soy Broth yielded the highest proportions of S-type cells (mean 88.7%), while BHI showed the lowest (mean 34.3%). CAMH and LB gave intermediate values (64.3% and 69.0%, respectively) (Fig. 4A). Temperature variation between 25°C and 42°C had minimal impact (Fig. 4B). In contrast, shaking speed had a strong effect on conversion efficiency: S-type cells consistently emerged at 120 rpm but were nearly absent at 180 rpm. Flask culture experiments confirmed this inverse correlation (Fig. 4C). Time-course analysis at 60 rpm showed that S-type cells began to increase notably around 12 hours (Fig. 4D), similar to tube culture observations.

**Fig. 4.**
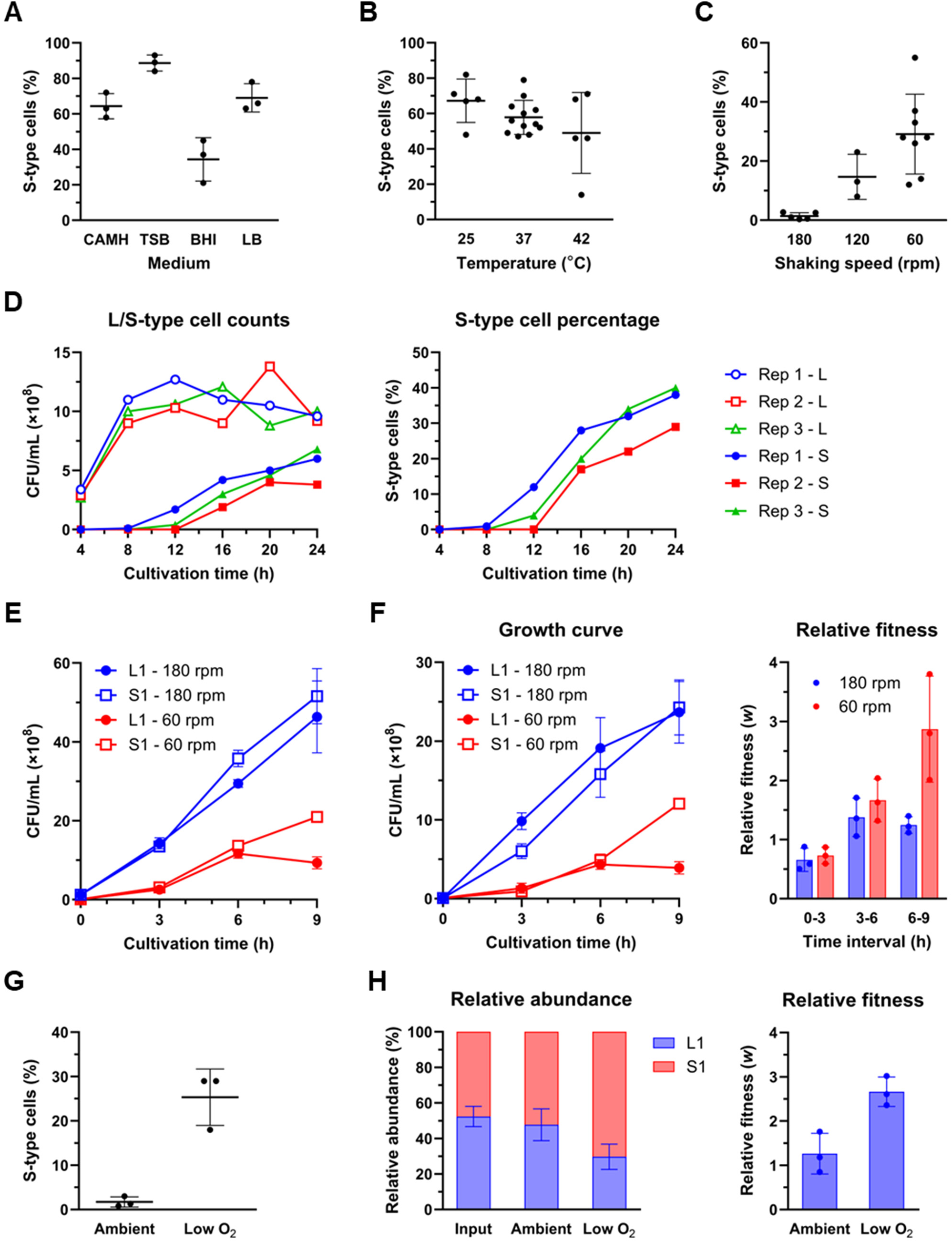
S-type cells have a fitness advantage under oxygen-limited conditions. (A) Effect of culture medium on L-to-S conversion. Single colonies of the L1 strain were inoculated into 3 mL of different media in 14-mL test tubes and incubated at 37°C with shaking at 120 rpm for 24 hours. The proportion of S-type cells was determined by plating and colony counting. Data represent means ± SD from three biological replicates (independent cultures initiated from single colonies), with individual data points shown. CAMH, cation-adjusted Mueller Hinton broth; TSB, Tryptic Soy Broth; BHI, Brain Heart Infusion broth; LB, Luria-Bertani broth. (B) Effect of temperature on L-to-S conversion. Single colonies of the L1 strain were inoculated into 3 mL of CAMH broth and incubated with shaking at 120 rpm at the indicated temperatures. Cultures were incubated for 24 hours at 37°C and 42°C, and for 48 hours at 25°C due to slower growth at the lower temperature. Data represent means ± SD from five biological replicates at 25°C and 42°C, and twelve biological replicates at 37°C (pooled data from the four independent experiments shown in Fig. 1C), with individual data points shown. (C) Effect of shaking speed on L-to-S conversion in Erlenmeyer flasks. Single colonies of the L1 strain were inoculated into 10 mL of CAMH broth in 50-mL Erlenmeyer flasks and incubated at 37°C for 24 hours at the indicated shaking speeds. Data represent means ± SD from five replicates at 180 rpm, three at 120 rpm, or eight at 60 rpm, with individual data points shown. (D) Time-course of S-type cell emergence at low shaking speed. Three independent L1 cultures (Rep 1 to 3) were grown in CAMH broth (10 mL in 50-mL flasks) at 37°C with shaking at 60 rpm. Samples were taken at indicated time points to determine CFUs of L-type and S-type cells. The left graph shows the CFUs for each type, and the right graph indicates the percentage of S-type cells in the population. (E) Growth curves of L1 and S1 strains in monoculture under different shaking conditions. Cultures were grown in CAMH broth (10 mL in 50-mL flasks) at 37°C with shaking at either 180 rpm or 60 rpm. Data represent means ± SD from three biological replicates. (F) Competition assay between L1 and S1 strains under different shaking conditions. L1 and S1 pWH1266 cells were mixed at a 1:1 ratio and cultivated in CAMH broth (10 mL in 50-mL flasks) at 37°C for 9 hours with shaking at either 180 rpm or 60 rpm. Left panel shows growth curves for both strains under the two conditions. Right panel shows the relative fitness (*w*) of S1 cells with respect to L1 cells, calculated for each time interval (0-3 h, 3-6 h, and 6-9 h). Data represent means ± SD from three biological replicates. (G) Effect of microaerophilic conditions on L-to-S conversion. Single colonies of the L1 strain were inoculated into 10 mL of CAMH broth in 50-mL flasks and incubated at 37°C with shaking at 180 rpm for 24 hours under ambient conditions or in sealed pouches with AnaeroPack MicroAero to create low-oxygen conditions. Data represent means ± SD from three biological replicates, with individual data points shown. (H) Competition assay under microaerophilic conditions. L1 and S1 pWH1266 cells were mixed at a 1:1 ratio and cultivated in CAMH broth at 37°C with shaking at 180 rpm for 9 hours under ambient or microaerophilic conditions (Low O_2_). Left panel shows the relative abundance of each strain at input and after 9 hours of cultivation. Right panel shows the relative fitness of S1 cells with respect to L1 cells under the two conditions. Data represent means ± SD from three biological replicates.

Having confirmed that S-type cells emerge and increase under low-shaking conditions, we hypothesized that these cells possess a growth advantage over L-type cells specifically under such conditions. To test this, we first compared the growth of L1 and S1 strains at 180 rpm and 60 rpm (Fig. 4E). At 180 rpm, both strains exhibited similar growth curves and continued proliferating throughout the 9-hour incubation. In contrast, at 60 rpm, L1 ceased growth after 6 hours, whereas S1 maintained steady proliferation, indicating a divergent response to reduced shaking. To further evaluate this growth advantage, we performed a competition assay using a 1:1 mixture of L1 and S1 strains at both shaking speeds (Fig. 4F). To distinguish the inoculated S1 cells from spontaneously emerging S-type cells, we used S1 carrying the pWH1266 plasmid, which confers resistance to ampicillin and tetracycline (16). At 180 rpm, the growth dynamics of L1 and S1 were largely comparable throughout the 9-hour cultivation. In contrast to the results at 180 rpm, and just as observed in the monoculture growth curve, L1 growth plateaued after 6 hours at 60 rpm, while S1 continued to proliferate, resulting in a markedly elevated relative fitness (*w*) for S1 during the 6–9 hour period (*w* = 2.87, SD 0.90). These results support the conclusion that S-type cells have a selective growth advantage under low-shaking conditions.

Since shaking speed affects aeration levels in bacterial cultures, we hypothesized that oxygen limitation promotes L-to-S conversion. To test this, we cultivated L1 strain under microaerophilic conditions using AnaeroPack MicroAero, which absorbs oxygen and generates carbon dioxide. Even at 180 rpm (where S-type emergence is normally suppressed), microaerophilic conditions considerably increased S-type proportions to 25.3% (SD 6.4) compared to 1.7% (SD 1.2) under ambient conditions (Fig. 4G). Competition assays further confirmed that oxygen limitation confers a selective advantage to S-type cells, with S1 exhibiting substantially higher relative fitness under microaerophilic conditions (*w* = 2.66, SD 0.33) than under ambient conditions (*w* = 1.26, SD 0.46) (Fig. 4H). These results demonstrate that oxygen limitation promotes L-to-S phenotypic conversion by providing S-type cells with a competitive advantage.

### L-to-S conversion mutations arise within individual L-type colonies

Because S-type cells became detectable as early as 12 hours after inoculation under low shaking conditions, we considered the possibility that L-to-S conversion might have occurred within the L-type colonies before inoculation. PCR screening using primers specific for IS*Aha2* insertions in *wzy* detected amplification products in multiple L1 colonies (Fig. 5A, B). Sequencing confirmed IS*Aha2* insertions at various positions and orientations (Fig. 5C), demonstrating mutation-carrying subpopulations within apparently homogeneous L-type colonies.

**Fig. 5.**
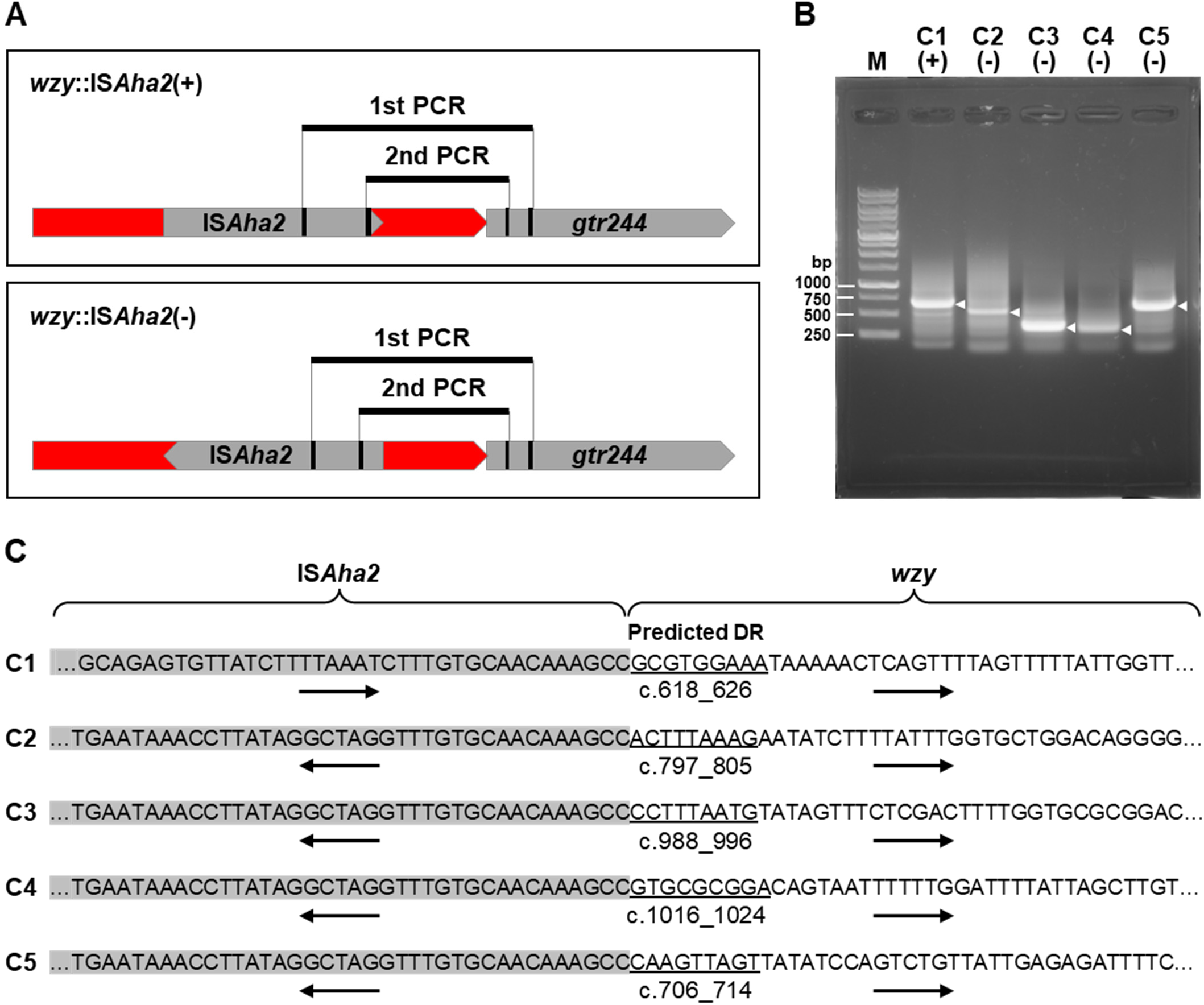
Detection of *wzy*::IS*Aha2* insertions within individual L-type colonies prior to liquid culture. (A) Schematic representation of the nested PCR strategy used to detect IS*Aha2* insertions in *wzy* in both forward (+) and reverse (−) orientations. PCR primers were designed to target sequences within IS*Aha2* and downstream of *wzy* (indicated by black segments), with a first-round PCR followed by a nested second-round PCR to increase specificity. Red arrows indicate *wzy*, and gray elements represent IS*Aha2* and the adjacent gene *gtr244*. (B) Agarose gel electrophoresis showing positive PCR products from nested PCR reactions. Eight individual L-type colonies were tested for IS*Aha2* insertions in both orientations (16 reactions in total), and only the five reactions yielding positive bands (from colonies C1 to C5) are shown. M indicates the molecular weight marker. (+) and (−) denote reactions designed to detect forward or reverse orientations of IS*Aha2* insertions, respectively. White arrowheads indicate bands that were excised and subjected to Sanger sequencing. (C) DNA sequence analysis of the PCR products shown in panel B, illustrating the junctions between IS*Aha2* and *wzy*. Gray highlighting indicates the terminal sequences of IS*Aha2*. Predicted direct repeat (DR) sequences are underlined, and the positions (e.g., c.618_626) refer to nucleotide coordinates within the *wzy* coding sequence corresponding to the DRs. Arrows below the sequences indicate the orientations of IS*Aha2* and *wzy*.

To understand why S-type colonies are not observed during routine subculturing on agar plates, we diluted and plated several L-type colonies. No S-type colonies were detected among over 10,000 screened colonies. These findings suggest that L-to-S conversion mutations arise spontaneously within colonies but remain at very low frequencies. The consistent emergence of S-type cells under oxygen-limited conditions is therefore likely driven by the selective expansion of these pre-existing mutants, although additional mutations during liquid culture cannot be excluded.

## Discussion

In this study, we characterized a clinical *A. baumannii* isolate that spontaneously gives rise to two morphologically and functionally distinct variants: an L type with a hypercapsule and an S type with reduced or absent capsule. The conversion from L to S occurred reproducibly under oxygen-limited conditions and was driven by multiple independent mutations within the *cps* gene cluster, including insertion sequence insertions and a single-nucleotide deletion. This phenotypic shift was associated with a trade-off between protective traits and environmental adaptability. While phenotypic switching has been previously reported in *A. baumannii* strain AB5075, that system is mediated by transcriptional regulation and is fully reversible, generating opaque and translucent colony types with differing virulence and resistance (17–21). In contrast, the L-to-S conversion observed in OCU_Ac16b is mutation-driven and unidirectional under our experimental conditions, reproducibly yielding S-type variants during liquid culture. These findings uncover a distinct, mutation-based mechanism of capsule variation that expands our understanding of how *A. baumannii* generates phenotypic diversity to adapt to environmental challenges.

Capsule biosynthesis loci in *A. baumannii* are frequently disrupted by ISs, as revealed by genome-wide surveys (22). Additionally, frameshift-like sequence anomalies have been identified in the *cps* locus in many genome assemblies, although their biological validity remains unclear (23). However, the functional and phenotypic consequences of such disruptions have remained largely unexplored. In this study, we provide experimental evidence that IS-mediated and point mutation-driven disruptions within the *cps* cluster reproducibly give rise to capsule-deficient variants. This demonstrates a dynamic mechanism of capsule switching that facilitates the rapid emergence of mutants with distinct survival advantages. Although our study focused on capsule loss, previous work using a different *A. baumannii* strain showed that a capsule-deficient state caused by an IS insertion in *itrA* can revert via scarless excision under specific conditions (22). These findings raise the possibility that *A. baumannii* may utilize both capsule loss and reversion as a reversible switching system to enhance environmental adaptability.

Similar mutation-driven capsule switching has also been observed in *Klebsiella pneumoniae*. Chiarelli et al. demonstrated that carbapenemase-producing *K. pneumoniae* clinical isolates frequently undergo mucoid-to-nonmucoid switching through diverse genetic events, including insertion sequence insertions (24). Unlike *A. baumannii* OCU_Ac16b, in which capsule-deficient variants do not visibly expand within colonies on agar plates, the nonmucoid mutants of *K. pneumoniae* rapidly grow within colonies to form distinguishable nonmucoid sectors. Accumulating evidence from both clinical and laboratory settings indicates that *K. pneumoniae* employs capsule switching as an integral strategy to survive in diverse environments (25–29). These parallel findings in *A. baumannii* and *K. pneumoniae* suggest that mutation-driven capsule switching may represent a shared adaptive strategy among encapsulated bacteria, facilitating phenotypic plasticity in response to environmental challenges. Our results highlight the potential impact of capsule switching on clinical diagnostics and infection outcomes. In routine clinical diagnostics, phenotypic or susceptibility testing is typically performed on a single colony isolated from a patient sample. If phenotypically distinct variants such as L- and S-type cells coexist within the same strain, this practice may lead to incomplete or misleading conclusions regarding the strain’s resistance or virulence profile. Beyond this diagnostic consideration, understanding the distinct characteristics of each variant is also important for predicting their behavior in clinical settings. For example, the desiccation tolerance of L-type cells may support prolonged survival on dry surfaces, facilitating environmental persistence and nosocomial transmission. Their enhanced resistance to serum killing may promote systemic dissemination once in the host. In contrast, S-type variants exhibit enhanced surface attachment and biofilm formation, favoring colonization of medical devices and persistence in chronic infections. Their superior growth under oxygen-limited conditions may confer a fitness advantage in hypoxic tissue environments such as necrotic wounds or inflamed lung tissue. These complementary traits suggest that reversible switching between L and S forms enables *A. baumannii* to exploit diverse niches and contributes to its persistence and success in clinical settings.

While our study provides novel insights into the mechanism and biological significance of capsule loss in *A. baumannii* OCU_Ac16b, several important questions remain unresolved. It is unclear whether the frequent mutations observed, particularly IS insertions, are specific to the *cps* cluster or whether similar mutations arise elsewhere in the genome but remain undetected— either because they impair growth and are counterselected, or because they confer no fitness advantage and thus fail to expand under the given culture conditions. The extent to which such mutations are reversible, either through IS excision or reversion mutations such as single-nucleotide duplications, also remains to be determined. In addition, the mechanisms underlying the fitness disadvantage of L-type cells under oxygen-limited conditions, as well as the increased β-lactam susceptibility associated with capsule loss, are not yet fully understood. Furthermore, it is essential to assess whether the observed phenomena are unique to OCU_Ac16b or represent a broader strategy among thickly capsulated clinical strains. Future studies should aim to comprehensively identify S-type mutations, assess their reversibility under relevant selection pressures, and compare gene expression profiles under aerobic and microaerobic conditions, in order to better understand the mechanisms governing capsule switching, including its regulatory control and responsiveness to environmental cues. Evaluating whether other thickly capsulated strains exhibit similar behavior under hypoxia will further clarify the generalizability and clinical relevance of our findings.

## Materials and Methods

### Bacterial strains and growth conditions

The *A. baumannii* strain OCU_Ac16b is a clinical isolate recovered from a patient at Osaka City University Hospital in 2015 (14). *A. baumannii* ATCC 19606^T^ served as a reference strain.

Cultivation was routinely performed in 14-mL round-bottom test tubes or 50-mL Erlenmeyer flasks containing 3 mL or 10 mL of CAMH broth, respectively. For experiments evaluating the proportion of S-type cells during or after 24-hour cultivation of L-type cells, the entire single colony was directly inoculated into the medium without prior adjustment of cell density. For experiments requiring OD_600_ or CFU normalization, such as competition assays and biofilm formation and adhesion assays, cells were scraped from agar plates, suspended in 1 mL of CAMH broth, and incubated at 37°C with shaking at 180 rpm for 1 hour prior to OD_600_ measurement, unless otherwise specified. We note that an OD_600_ of 1.0 corresponds to approximately 1 × 10^9^ CFU/mL for both L-type and S-type strains under our experimental conditions.

For microaerophilic cultivation, flasks were placed in sealed plastic pouches together with an AnaeroPack MicroAero (Mitsubishi Gas Chemical Co., Inc.), which absorbs oxygen and generates carbon dioxide to create low-oxygen conditions. The pouches were then incubated at 37°C with shaking at 180 rpm.

### Competition assay

To evaluate the relative fitness of S-type cells compared to L-type cells, L1 and S1 pWH1266 cells were preincubated in CAMH broth for OD_600_ adjustment, as described in the *Bacterial strains and growth conditions* section. Both strains were then inoculated into 10 mL of CAMH broth in 50-mL Erlenmeyer flasks at a 1:1 ratio, with a final concentration of approximately 1 × 10^7^ CFU/mL for each strain. Cultures were incubated at 37°C for 9 hours under different conditions: with shaking at either 180 rpm or 60 rpm, or at 180 rpm under microaerophilic conditions using AnaeroPack MicroAero. After incubation, bacterial suspensions were appropriately diluted and plated on CAMH agar with or without tetracycline (10 µg/mL) and ampicillin (100 µg/mL) to determine the proportion of S-type colonies.

Relative fitness (*w*) of S-type cells was calculated using the equation, *w* = [*x*_2_(1 − *x*_1_)] / [*x*_1_(1 − *x*_2_)], where *x*_1_ is the initial frequency of S-type cells and *x*_2_ is the final frequency (30).

### Genome sequencing and mutation identification

Cells grown overnight on CAMH agar plates were harvested and resuspended in PBS. Genomic DNA was extracted using Genomic-tip 100/G (Qiagen, Hilden, Germany) according to the manufacturer’s instructions.

Whole-genome sequencing of L1 and S1 was performed using both long-read (PacBio Sequel IIe) and short-read (Illumina HiSeq X) platforms. HiFi reads (∼250 Mb per sample) were generated using the SMRTbell Express Template Prep Kit 2.0 (Pacific Biosciences, Menlo Park, CA, USA). Paired-end 150-bp Illumina reads were prepared with the NEBNext Ultra II DNA Library Prep Kit (New England Biolabs, Ipswich, MA, USA). For strains S2–S5, sequencing was performed on the Illumina NovaSeq 6000 platform using the NEBNext Ultra II kit.

HiFi reads were filtered with Filtlong v0.2.1 (minimum length, 2,000 bp; minimum quality score, 20) to retain the top 90%, targeting 230 Mb. The N50 values of L1 and S1 were 14,951 bp and 13,931 bp, respectively. Illumina reads were trimmed using fastp v0.23.1 (31). For L1, filtered HiFi reads were assembled *de novo* into four circular contigs with Flye v2.9.2 (32), then polished twice with Pilon v1.24 (33) using BAM files from both PacBio and Illumina reads. For S1, HiFi reads were mapped to the L1 genome using Minimap2 (34), and a consensus sequence was polished with Pilon using Illumina reads to yield the final assembly.

To identify mutations responsible for phenotypic conversion, S-type genomes were compared with L1. For S1, structural and point mutations were detected by aligning the assembled genome with L1 using breseq (35). For S2–S5, quality-filtered Illumina reads were used for both *de novo* assembly and reference-based variant analysis. Contigs spanning the *cps* cluster were compared with the L1 reference, and genome-wide variants were identified by read mapping, variant calling, and manual curation.

During genome analysis of OCU_Ac16b (L1 and S1), we identified three plasmids: pOCU_Ac16a_1, pOCU_Ac16a_3, and an additional plasmid, pOCU_Ac16a_4, which had not been detected in our previous analysis of the clonally related strain OCU_Ac16a. Re- examination of the original sequencing data revealed that pOCU_Ac16a_4 had been present in Ac16a but was missed during the initial assembly.

### Antimicrobial susceptibility tests

Antimicrobial susceptibility testing was performed using Etest strips (bioMérieux, Marcy-l’Étoile, France) in accordance with the guidelines of the Clinical and Laboratory Standards Institute.

### Serum resistance assay

Human blood was collected from three healthy donors (all coauthors of this study), combined, and used to prepare pooled serum. The blood was left at room temperature for 2 hours to allow clotting, then centrifuged. The resulting serum was aliquoted and stored at −80°C until use. Heat-inactivated serum was prepared by incubation at 56°C for 30 minutes.

For the serum resistance assay, pooled serum was diluted with PBS to final concentrations of 2.5% and 10%. Cells of the tested strains were inoculated from single colonies into CAMH broth, cultured for 6 hours, and then washed twice with PBS. The resulting cell suspension was adjusted to an OD_600_ of 1.0. A 10-μL aliquot of this suspension (∼1 × 10⁷ CFU) was added to 200 μL of diluted serum in a 96-well plate and incubated at 37°C with shaking at 100 rpm for 2 hours. After incubation, the mixtures were serially diluted and plated to determine CFU, and survival rates were calculated.

### Biofilm formation and adhesion assays

For biofilm formation assays, cells preincubated in CAMH broth for OD_600_ adjustment, as described in the *Bacterial strains and growth conditions* section, were inoculated into 1 mL of BHI medium in each well of 24-well polystyrene plates at an initial OD_600_ of 0.01. The plates were incubated under static conditions at 37°C for 48 hours. After incubation, the culture medium was discarded, and the wells were washed once with PBS and air-dried completely. Biofilms were stained with 2 mL of 0.1% crystal violet for 20 minutes, washed three times with PBS, and air-dried. The dye was then solubilized with 2 mL of 95% ethanol, and the absorbance was measured at 590 nm.

For adhesion assays, cells were inoculated into 1 mL of CAMH broth per well in 24-well plates at an initial OD_600_ of 0.1 and incubated at 37°C with shaking at 100 rpm for 2 hours. After incubation, the medium was discarded, and the wells were gently washed twice with PBS to remove planktonic cells. Attached cells were stained and quantified as described for the biofilm assay.

### Desiccation tolerance assay

Cells were pre-cultured in CAMH broth at 37°C with shaking (180 rpm) for 3 hours, washed three times with distilled water, and resuspended to an OD_600_ of 0.1. Ten microliters of suspension was dispensed into 96-well plates and incubated at 36°C under 13 ± 3% relative humidity. At designated time points (1, 2, 3, 4, 5, 12, or 21 days), cells were rehydrated with 100 μL distilled water and incubated at 36°C for 30 minutes. After resuspension by pipetting, samples were serially diluted and plated to determine CFU.

### PCR screening for *wzy*::IS*Aha2* insertions in individual L-type colonies

To detect IS*Aha2* insertions in *wzy* within single colonies, nested PCR was performed as illustrated in Fig. 5A. Colonies were obtained by directly streaking glycerol stocks onto agar plates and incubating overnight (∼18 hours) at 37°C. Entire well-isolated colonies were resuspended in 30 µL of sterile water and heat-treated at 99°C for 15 minutes. Each sample was then split into two 15-µL aliquots, both of which were used in separate first-round PCR reactions targeting opposite orientations of IS*Aha2* insertions. First-round PCR was conducted using MightyAmp DNA Polymerase Ver.3 (Takara Bio, Shiga, Japan), followed by nested PCR using Ex Taq polymerase (Takara Bio). Primer sequences are listed in Table 1. PCR products were gel-analyzed, purified, and Sanger sequenced to confirm insertion and determine the site.

### Use of generative AI tools

During the manuscript preparation process, generative AI tools (ChatGPT by OpenAI and Claude by Anthropic) were used in a limited capacity to assist with English language editing and phrasing suggestions. All scientific content, data interpretation, and conclusions are solely the work of the authors.

## Data availability

The complete sequences of the chromosome and three plasmids (pOCU_Ac16a_1, pOCU_Ac16a_3, and pOCU_Ac16a_4) from strains L1 and S1 will be deposited in the DDBJ/EMBL/GenBank databases upon journal publication. The DDBJ entry for the strain OCU_Ac16a (AP023077–AP023080) will be updated to include pOCU_Ac16a_4. Raw sequencing data (PacBio and Illumina) for strains L1 and S1, as well as Illumina data for S2–S5, will be deposited in the DDBJ Sequence Read Archive upon journal publication.

## Acknowledgments

We are grateful to JST SPRING (Grant Number JPMJSP2139) for providing financial support and career development training to C.S. and A.S. C.S. was additionally supported by the Osaka Metropolitan University Research Project Scholarship (2021–2025) and the Yang DaPeng Scholarship (2021–2023). This research was also supported by the Japan Agency for Medical Research and Development (AMED) (Grant Numbers JP25fk0108665, JP25fk0108683, JP25fk0108712, JP25wm0225029, and JP25gm1610003 to M.S.) and by JSPS KAKENHI (Grant Numbers 23K06556 to M.S., 24K11636 to Y.K., and 22K07072 to K.O.).

## Author contributions

**Chaogetu Saren:** Conceptualization; Formal analysis; Funding acquisition; Investigation; Methodology; Resources; Visualization; Writing – Original Draft; Writing – Review & Editing

**Ken-Ichi Oinuma:** Conceptualization; Data curation; Formal analysis; Funding acquisition; Methodology; Project administration; Supervision; Writing – Original Draft; Writing – Review & Editing

**Taishi Tsubouchi:** Data curation; Formal analysis; Writing – Review & Editing

**Arata Sakiyama:** Formal analysis; Investigation; Writing – Review & Editing

**Masato Suzuki:** Conceptualization; Data curation; Funding acquisition; Methodology; Writing – Review & Editing

**Mamiko Niki:** Methodology; Resources; Supervision

**Yukihiro Kaneko:** Conceptualization; Funding acquisition; Investigation; Resources; Supervision; Writing – Review & Editing

